# The *Staphylococcus aureus* LXG-domain toxins EsxX and SAR0287 do not promote virulence in a zebrafish larval infection model

**DOI:** 10.1101/2025.10.04.680436

**Authors:** Fatima Ulhuq, Amy Tooke, Chriselle Mendonca, Guillermina Casabona, Johann Habersetzer, Yaping Yang, Margarida C. Gomes, Felicity Alcock, Serge Mostowy, Tracy Palmer

**Affiliations:** Newcastle University Biosciences Institute, Newcastle University, Newcastle upon Tyne, NE2 4HH, UK; School of Life Sciences, University of Dundee, Dundee DD1 5EH, UK; Department of Infection Biology, London School of Hygiene & Tropical Medicine, WC1E 7HT, UK; School of Life Sciences, University of Warwick, CV4 7AL, UK

**Keywords:** type VII secretion system, *Staphylococcus aureus*, membrane-depolarising toxin, toxin-immunity, interbacterial competition, zebrafish

## Abstract

The *Staphylococcus aureus* type VIIb secretion system (T7SSb) is a multiprotein secretion system that secretes toxins with antibacterial activity, but which is also required for full virulence in animal models of infection. *S. aureus* strains carry one of four T7SSb locus types, named *essC1* to *essC4*, each of which encodes a characteristic LXG-family substrate at the T7SS locus. In *essC2* strains this LXG-domain protein is EsxX, which has a glycine zipper sequence in its C-terminus and has potent antibacterial membrane-depolarising activity. In this work we recognise conserved features of the *essC2* and *essC3* systems, identifying the LXG protein SAR0287 as structurally and functionally similar to EsxX. Using a zebrafish larval hindbrain ventricle infection model we demonstrate that the T7SSb of *essC2* and *essC3* representative strains contribute to bacterial replication and zebrafish mortality. However, there is no significant loss of virulence in the model system if EsxX or SAR0287 are absent. These findings indicate there is no discernible role for either toxin in this virulence model.

## Introduction

*Staphylococcus aureus* is an opportunistic pathogen of humans and economically-important livestock such as pigs and cattle. In humans, the anterior nares is the most common carriage site with 20-30% of individuals estimated as asymptomatic carriers (1). *S. aureus* is easily spread via skin-to-skin contact and can breach the epithelial barrier to cause skin and soft tissue infections, pneumonia, sepsis, or a variety of other clinically important infections. Methicillin-resistant *S. aureus* (MRSA) is a major cause of infections worldwide and is becoming increasingly difficult to treat due to emerging resistance to all antibiotic classes in MRSA lineages (2).

All *S. aureus* strains encode a single copy of the type VIIb protein secretion system (T7SSb), which is important for both virulence and bacterial antagonism. The overall contribution of the *S. aureus* T7SSb to virulence is well established (3-10), however the precise pathogenic roles of specific secreted effectors are less clear. Two classes of secreted substrate have been described: small helical hairpin proteins with similarity to the WXG100 family, and polymorphic antibacterial toxins (11). The latter are multi-domain proteins which in addition to the toxin domain have an N-terminal LXG domain, or a structurally similar C-terminal reverse-LXG domain (9, 12). These are helical domains that target the toxin to the T7SSb, in concert with cognate WXG100-like partner proteins, termed Laps (for **L**XG **a**ccessory **p**roteins) which bind to the LXG domain and are co-secreted with the toxin (12-15). In addition to these toxin-specific WXG100-like secretion factors, the core T7SSb component, EsxA, is a WXG100 family protein found in all strains which is secreted as a homodimer (16, 17).

The *S. aureus* T7SSb is encoded by the *ess* locus, of which there are four sequence variants named *essC1* to *essC4* after the sequence variant of the key T7SS core component EssC that they encode (16). Other conserved components of the secretion machinery are encoded upstream of *essC*, while downstream is an assortment of variant-specific effectors, accessory secretion factors and immunity genes. Each *ess* locus carries a single variant-specific LXG domain-encoding gene, and strains also encode additional conserved toxin substrates such as TspA and TslA at other loci (9, 10, 18). The *essC1* locus encodes an LXG-nuclease toxin, EsaD, with an established role in bacterial antagonism (10, 19). *essC2* loci encode an LXG-glycine zipper toxin, EsxX, which depolarises the bacterial cytoplasmic membrane (20), while the *essC3-* and *essC4-*specific LXG family substrates have yet to be characterised. In a prior study, deletion of *esxX* in an ST398 clinical isolate reduced bacterial replication and abscess formation in mice and resulted in a decrease in lysis of human neutrophils *in vitro* (21). Thus, EsxX might function in both interbacterial and host interactions. We have previously developed a zebrafish larval infection model using the sterile hindbrain ventricle as an infection site to study *S. aureus*-host interactions in the absence of confounding interbacterial interactions. Using this model we determined that the conserved substrate TspA, but not the variant-specific EsaD toxin, contributes to zebrafish mortality (10). In this work we use the same model to investigate the requirement for EsxX and its *essC3* equivalent SAR0287 in *S. aureus* virulence.

## Methods

### Construction of bacterial strains and plasmids

*S. aureus* strains used in this work are listed in Table 1. Strain 10.1252.X::apra was constructed using the pIMAY system (22). Plasmids are listed in Table S1. Plasmid pIMAY-apra-ins, carries an apramycin resistance gene (*aac(3)-IVa*) under the control of the constitutive *rpsF* promoter, with flanking sequences corresponding to an intergenic region downstream of *SAPIG0009*. The cassette carrying the flanking regions, promoter and resistance gene was amplified as four fragments, joined by overlap PCR then cloned into the EcoRI-SacI sites of pIMAY (22). Plasmid pIMAY-GFP-ins encodes GFP under control of the *S. aureus sarA* promoter and a strong terminator (amplified from pTH100; reference 23). It is flanked by DNA encoding fragments of *SAPIG0102* and *SAPIG0103* and designed to be inserted in the intergenic region. The flanking regions and *gfp* cassette were amplified as three fragments, joined by overlap PCR and cloned into the EcoRI site of pIMAY. Chromosomal deletions were constructed by allelic exchange using plasmid pIMAY carrying the flanking regions (∼500 bp each) of the deleted region. pIMAY_SAPIG0305 (EcoRV), pIMAY_SAR0287 (EcoRI-BamHI) and pIMAY-ess-M252 (KpnI-SalI) were created by amplifying the two flanking regions, joining them together by overlap PCR, then restriction cloning the ∼1kb fragment into the indicated sites of pIMAY. For expression of SAR0287 fragments in *E. coli*, pBAD18-cm was used (24). All primers used for strain and plasmid construction are listed in Table S2. *E. coli* strains DH5α (*F*^-^φ*80dlacZM15* (*lacZYA-argF*)*U169 deoR recA1 endA1 hsdR17(r k*^*-*^, *mk*^*+*^*) phoA supE44 thi-1 gyrA96 relA1* λ^-^) and JM110 (*rpsL* (Strr) *thr leu thi-1 lacY galK galT ara tonA tsx dam dcm supE44* Δ(*lac-proAB*) [F′*traD36 proAB lacI*q*Z*Δ*M15*]) were used for cloning and preparation of plasmids for electroporation into *S. aureus*, respectively, while MG1655 (F-lambda- *ilvG*- *rfb*-50 *rph*-1) was used to express proteins from pBAD plasmids. *S. aureus* strains were cultured in tryptic soy broth (TSB, Oxoid) containing either 25 μg/ml apramycin (for 10.1252.X derivative strains that carried an apramycin resistance gene) or 5 μg/ml erythromycin (for MRSA252 and derivative strains). *E. coli* was cultured in Luria broth (LB, Melford).

**Table 1.**
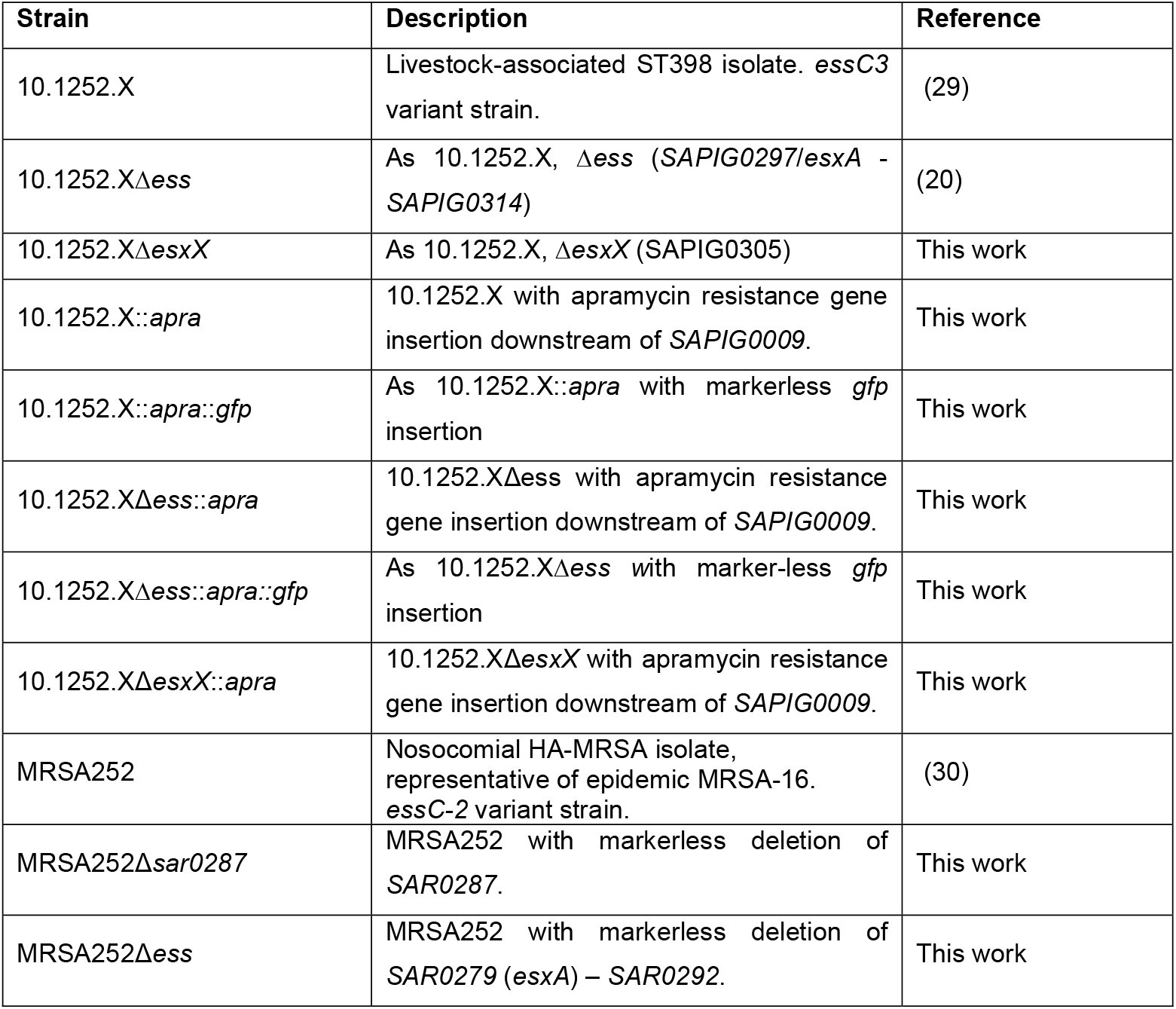
*S. aureus* strains used in this study.

### Zebrafish larva infection

All experiments using zebrafish were carried out as previously described (10). Briefly, embryos were obtained from wild-type (AB strain) zebrafish. Hindbrain ventricle infections were carried out at 3 days post-fertilization (dpf) and incubated at 33 °C, following injection of 1-2 nl of bacterial suspension containing the required dose. At the indicated times, larvae were killed in tricaine, lysed with Triton X-100 and homogenized mechanically. Larval homogenates were serially diluted and plated onto TSB agar for CFU enumeration. Competition experiments were carried out as previously (10). 3 day post-fertilization zebrafish larvae anaesthetised with tricaine were injected in the hindbrain ventricle with 1-2 nl bacterial suspension containing 6000 CFU of each strain (a 1:1 ratio). After removal of injured larvae, they were washed in E2 buffer to remove tricaine. Larvae were incubated at 33 °C. At indicated timepoints, larvae were selected without bias (first 8 living larvae observed) and mechanically homogenised in E2, before being serially diluted in PBS and plated out onto TSA and incubated at 37 °C overnight to determine CFU per larva. Animal experiments were performed according to the Animals (Scientific Procedures) Act 1986 and approved by the UK Home Office (Project licenses: PPLP84A89400 and P4E664E3C). Competition indices were calculated as follows where values for “prey” and “attacker” are the mean CFU for the group at the indicated timepoint (n):

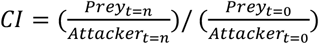

Where t=0 is 0 hours post infection (hpi) and t = n is 2, 4 or 6 hpi. Fig. 4D shows the mean of the indices for each timepoint for the 2 biological replicates.

## Results

### Contribution of the ST398 T7SS and EsxX to virulence in a zebrafish larval infection model

*esxX* is present at the *essC2* locus in *S. aureus* ST398 lineages and has been reported to promote virulence in mice (21) and to depolarise the bacterial cytoplasmic membrane (20). Using a previously developed zebrafish larval infection model in which the sterile hindbrain ventricle is inoculated with bacteria (10, 25) we first assessed virulence of an ST398 isolate. Survival of zebrafish larvae was monitored following inoculation of the hindbrain ventricle with the *S. aureus* ST398 isolate 10.1252.X at 3 days post-fertilization (dpf) (Fig. 1A,B). Dose-dependent mortality was observed at 24 h post-infection (hpi) with ∼80% of zebrafish larvae surviving a low dose of *S. aureus* (∼900 CFU, ‘dose 1’) and only ∼40% surviving a higher dose (∼7.5 x 10^3^CFU, ‘dose 2’). Using the higher dose inoculum, we next investigated whether the entire *ess* gene cluster, or EsxX specifically, contributes to zebrafish mortality. We compared survival of larvae injected with wild-type 10.1252.X to those injected with either the single deletion strain 10.1252.XΔ*esxX*, or strain 10.1252.XΔ*ess* lacking all 18 genes at the *ess* locus (from *SAPIG0297* to *SAPIG0314*). Mortality was significantly reduced at 24 hpi for larvae infected with the 10.1252.X*Δess* strain compared to the wild-type, suggesting a role for the T7SS or its effectors in virulence (Fig. 1C). In agreement, total bacterial counts of infected larvae revealed that there was a significant decrease in recovery of the Δ*ess* strain compared to the wild-type at 4 hpi suggesting that bacteria lacking the T7SS proliferate less well *in vivo* (Fig. 1D). This is in agreement with a previous report noting decreased virulence of an ST398 T7SS mutant strain in murine skin and blood infection models (8). However, no significant difference was observed for either larval survival or bacterial recovery counts between the wild-type and Δ*esxX* strains (Fig. 1C,D). This contrasts with the study of Dai *et al*. (21) who reported a reduced virulence phenotype in mouse models when *esxX* was deleted. Control experiments demonstrated that neither mutant exhibited a growth defect in standard laboratory culture (Fig. S1). We therefore conclude that the observed contribution of the T7SS to virulence in the zebrafish larval model is not driven by EsxX secretion.

### Conserved features of the *S. aureus* T7SS *essC2* and *essC3* variants

Of the four *ess* locus variant types found in *S. aureus*, the two variants that are most similar are *essC2* and *essC3* (Fig. 2A). A single LXG domain-containing protein, SAR0287, is encoded at the variable region of the *essC3* locus, and structural homology searches indicate that like EsxX, SAR0287 is an LXG toxin with a probable channel-forming C-terminal domain. Indeed, the C-terminal regions of EsxX and SAR0287 share 45% sequence identity, including conservation of an extended glycine zipper motif which is required for toxicity (20) (Fig. 2B). Homologues of the EsxX secretion factors (LapX3, LapX4) and immunity proteins (ExiA, ExiB, ExiC and ExiD) are also encoded at the *SAR0287* locus with conserved synteny (Fig. 2C). This strongly suggests that, like EsxX, SAR0287 is a glycine zipper toxin with antibacterial activity. In agreement with this, the C-terminal glycine zipper domain of SAR0287 was highly toxic to *E. coli* even when expression of the protein was repressed in the presence of glucose (Fig. 2D). Moreover, all our attempts to clone *SAR0287* for regulated expression in *S. aureus* were unsuccessful, consistent with the protein having antibacterial activity.

**Figure 1.**
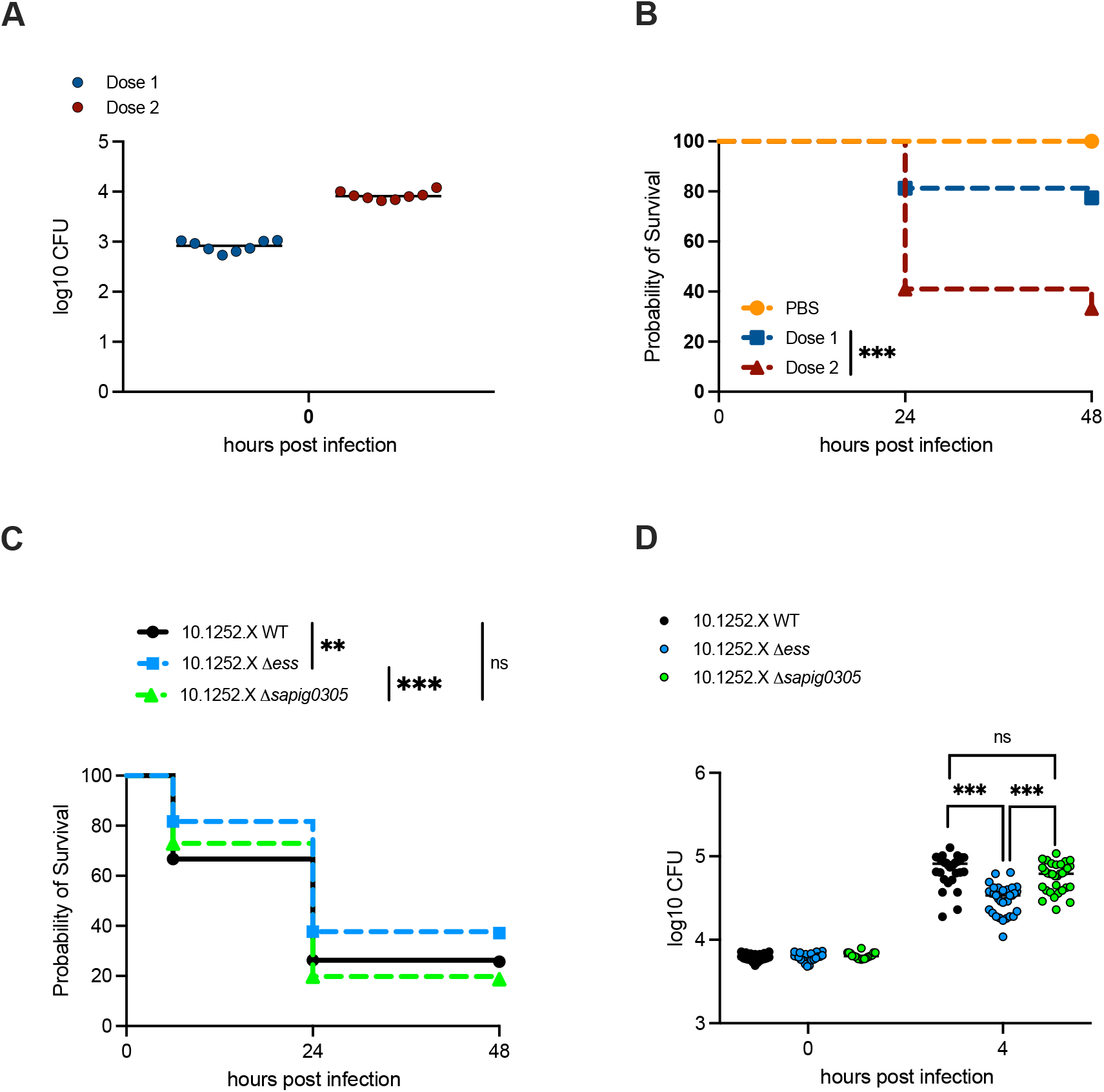
Contribution of EsxX to virulence in a zebrafish larva infection model. A. Enumeration of bacteria recovered at 0 hpi from zebrafish embryos infected with isolate 10.1252.X::*apra*. Data are pooled from 2 independent experiments where 4 larvae were sacrificed at 0 hours to determine the input dose. Circles represent individual larvae, horizontal bars show the mean. B. Survival curves of zebrafish larvae injected in the hindbrain ventricle with 10.1252.X::*apra*. Zebrafish were injected at 3 dpf with a low (∼900 CFU, dose 1) or high (∼7.5 x 10^3^ CFU, dose 2) dose of 10.1252.X::*apra*, incubated at 33 °C and monitored for 48 hpi. Data are pooled from two independent experiments (n=36-44 larvae per experiment). Results are plotted as Kaplan-Meier survival curves and the p value between conditions was determined by log-rank Mantel-Cox test. **** p<0.0001. Additional embryos were inoculated for bacterial enumeration at 0h, shown in (A). C. Survival curves as in (B) but with the indicated strains (all apra^r^) at dose 2. Data are pooled from four independent experiments (n=38-55 larvae per strain per experiment). Results are plotted as Kaplan-Meier survival curves and the p value between conditions was determined by log-rank Mantel-Cox test. ** p<0.01, ns, not significant. Additional embryos were inoculated for bacterial enumeration at 0h and 4h, shown in (D). D. Enumeration of recovered bacteria at the indicated timepoints from zebrafish embryos infected with the indicated strains during the survival experiment shown in (C). Pooled data from 4 independent experiments where 5-8 larvae were sacrificed per timepoint. Only larvae having survived the infection were included. Circles represent individual larvae, horizontal bars represent the mean. Significance was tested using an unpaired t test. **** p<0.0001, ns, not significant.

**Figure 2.**
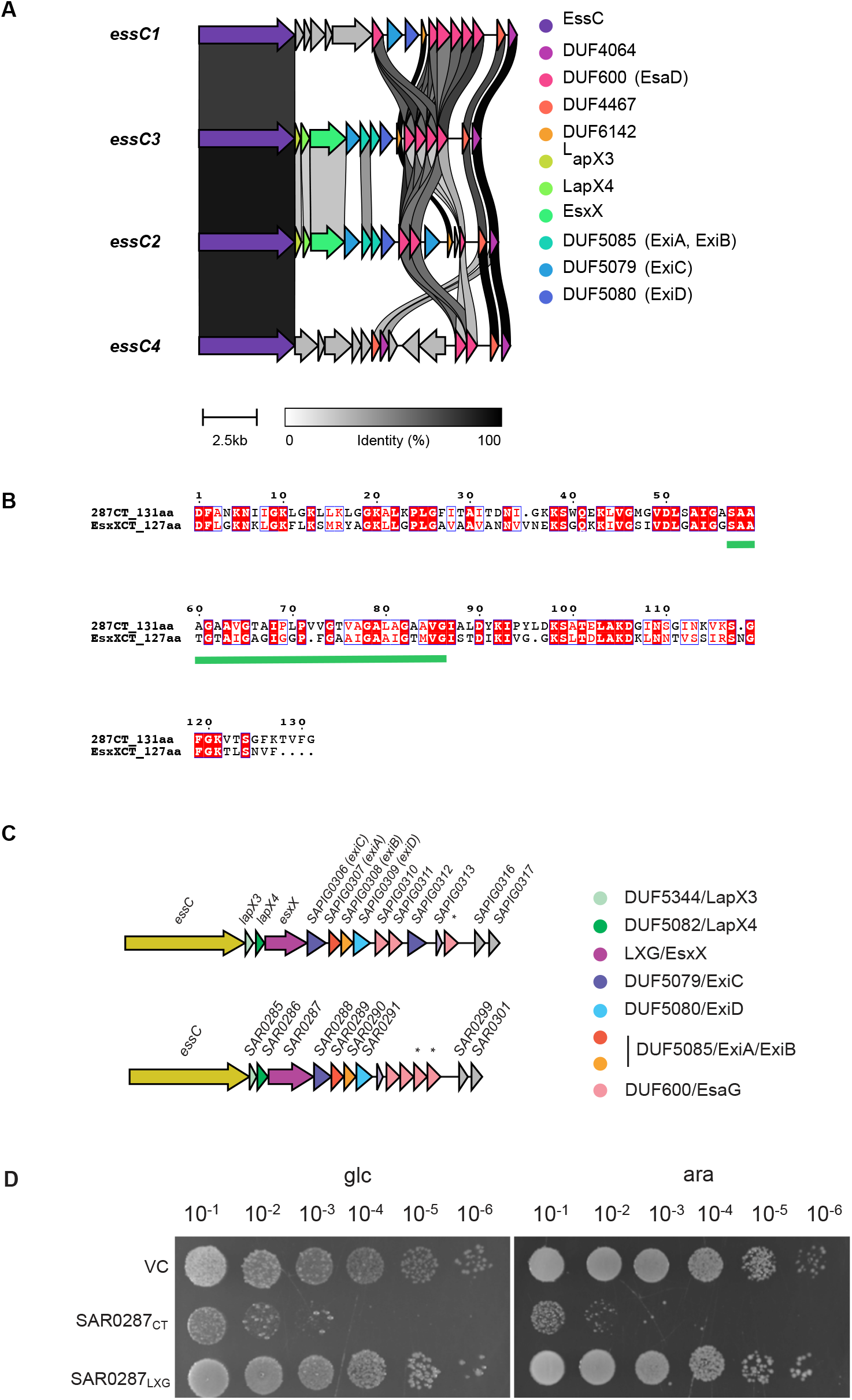
Conserved genes in the *ess* loci of *S. aureus* T7SS *essC2* and *essC3* variants. A. Comparison of the gene clusters downstream of *essC* in strains representing each of the four *S. aureus essC* variant locus types (*essC1*, NCTC8325, accession CP000253; *essC3*, MRSA252, accession BX571856; *essC2*, ST398, accession NC_017333; and *essC4*, HO 5096 0412, accession HE681097). Homologous genes are colour coded and connected by grey links, shaded according to percent identity. Figure was created using Clinker (32). B. Protein sequence alignment of the C-terminal 131 residues of SAR0287 with the C-terminal 127 residues of EsxX. An extended glycine zipper motif conserved in both sequences is indicated with a green bar. C. Schematic diagram of the genes downstream of *essC* in ST398 and MRSA252, colour-coded by protein family. *pseudogene. D. Overnight cultures of *E. coli* strain MG1655 carrying empty pBAD18-cm (VC) or pBAD18-cm encoding the SAR0287 C-terminal (SAR0287_CT_; residues 315 – 556) or N-terminal (SAR0287_LXG_; residues 1-314) regions were serially diluted and spotted onto LB plates containing either 0.5% D-glucose or 0.02% L-arabinose, as indicated. Plates were incubated overnight at 37°C.

### Contribution of the MRSA252 T7SS and SAR0287 to virulence in a zebrafish infection model

To investigate potential roles of the MRSA252 T7SS and SAR0287 in bacterial virulence, we deleted *SAR0287* or the entire *ess* locus (from *SAR0279*/*esxA* – *SAR0292*) from this strain and assessed the effect of these mutations on zebrafish larval mortality, as described above for 10.1252.X. Figure 3B shows that, like 10.1252.X, MRSA252 also causes larval mortality in a dose-dependent manner, with ∼80% survival at a low dose of ∼8 x 10^3^ CFU (dose 1) and ∼40% survival when injected with a higher dose of ∼2 x 10^4^ CFU (dose 2), at 48 hpi (Fig. 3A,B). Comparing the wild-type strain with the MRSA252Δ*ess* and MRSA252Δ*SAR0287* deletion strains, again at the higher dose inoculum, reveals a similar picture to that observed for 10.1252.X; when inoculated with the Δ*ess* strain, zebrafish mortality was significantly reduced at 48 hpi compared to wild-type, and bacterial cell counts were also reduced at 20 hpi for the Δ*ess* mutant (Fig. 3C,D). By contrast, deletion of the SAR0287 toxin gene alone, did not significantly affect either mortality or bacterial cell counts relative to the wild-type strain (Fig. 3C,D). Thus, while there is a significant reduced virulence phenotype associated with deletion of the T7SS/*ess* loci in both the *essC2* strain 10.1252.X and the *essC3* strain MRSA252, this decreased virulence does not appear to result from loss of EsxX or SAR0287 secretion.

**Figure 3.**
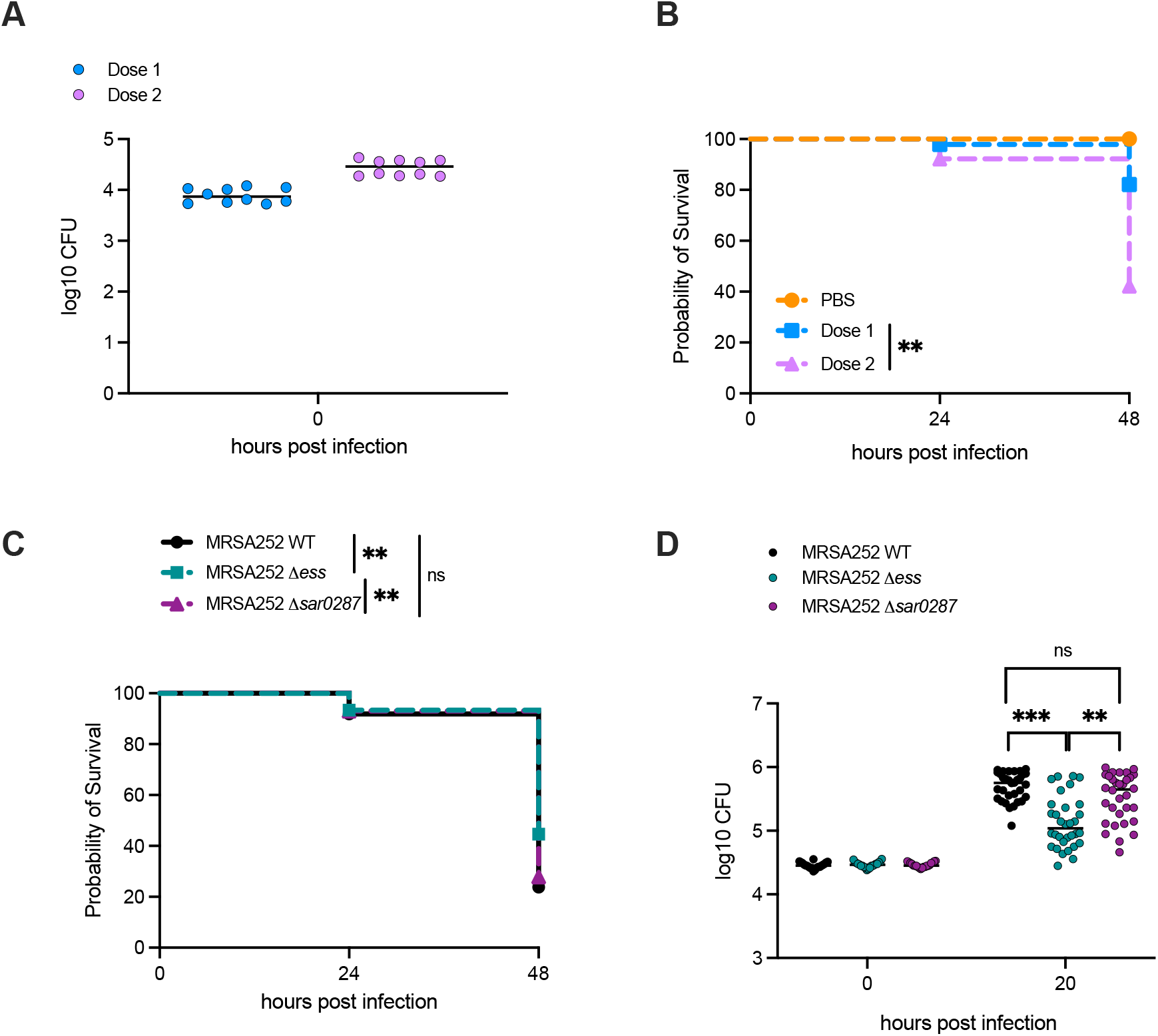
Contribution of SAR0287 to virulence in a zebrafish larva infection model. A. Enumeration of bacteria recovered at 0 hpi from zebrafish embryos infected with isolate MRSA252. Data are pooled from 2 independent experiments where 5 larvae were sacrificed at 0 hours to determine the input dose. Circles represent individual larvae, horizontal bars show the mean. B. Survival curves of zebrafish larvae injected in the hindbrain ventricle with WT MRSA252. Zebrafish were injected at 3 dpf with a low (∼900 CFU, dose 1) or high (∼7.5 x 10^3^ CFU, dose 2) dose of MRSA252, incubated at 33 °C and monitored for 48 hpi. Data are pooled from two independent experiments (n=36-44 larvae per experiment). Results are plotted as Kaplan-Meier survival curves and the p value between conditions was determined by log-rank Mantel-Cox test. *** p<0.001. Additional embryos were inoculated for bacterial enumeration at 0h, shown in (A). C. Survival curves as in (B) but with the indicated strains at dose 2. Data are pooled from four independent experiments (n=38-55 larvae per strain per experiment). Results are plotted as Kaplan-Meier survival curves and the p value between conditions was determined by log-rank Mantel-Cox test. *** p<0.001, ns, not significant. Additional embryos were inoculated for bacterial enumeration at 0h and 20h, shown in (D). D. Enumeration of recovered bacteria at the indicated timepoints from zebrafish embryos infected with the indicated strains during the survival experiment shown in (C). Pooled data from 4 independent experiments where 5-8 larvae were sacrificed per timepoint. Only larvae having survived the infection were included. Circles represent individual larvae, horizontal bars represent the mean. Significance was tested using an unpaired t test. **** p<0.0001, ns, not significant.

### Assessing EsxX-dependent intrastrain competition in the larval hindbrain

We have reported that the sterile larval hindbrain can be used to assess competition between pairs of *S. aureus* strains. In those experiments the nuclease toxin EsaD from the *essC1* strain COL was able to inhibit growth of a different *essC1* strain, RN6390, provided that the genes for the EsaD immunity protein, EsaG and its homologues were deleted (10). To assess EsxX-dependent intrastrain competition, we competed 10.1252.X against a GFP-expressing but otherwise isogenic 10.1252.X strain, or GFP-expressing 10.1252.X lacking all genes at the T7SS locus (10.1252.XΔ*ess*; Fig 4A), including *exiC* and *exiD* which provide immunity against secreted EsxX (20). While we had previously assessed bacterial survival by measuring colony counts after 15 and 24 hours (10), the high lethality of 10.1252.X resulted in death of the majority of larvae at these timepoints, so we sampled colony counts 2 - 6 hours post infection. As shown in Fig 4B,C there was no difference in recovered counts between the ‘attacker’ strain (10.1252.X) and either of the GFP-expressing ‘prey’ strains. In agreement with this, the competitive indices were close to 1 for all strain combinations and timepoints (Fig. 4D). These findings indicate that the strain lacking an active T7SS and the EsxX immunity genes was not disadvantaged in competition with the wild-type and thus there is no evidence of intrastrain warfare under these conditions.

**Figure 4.**
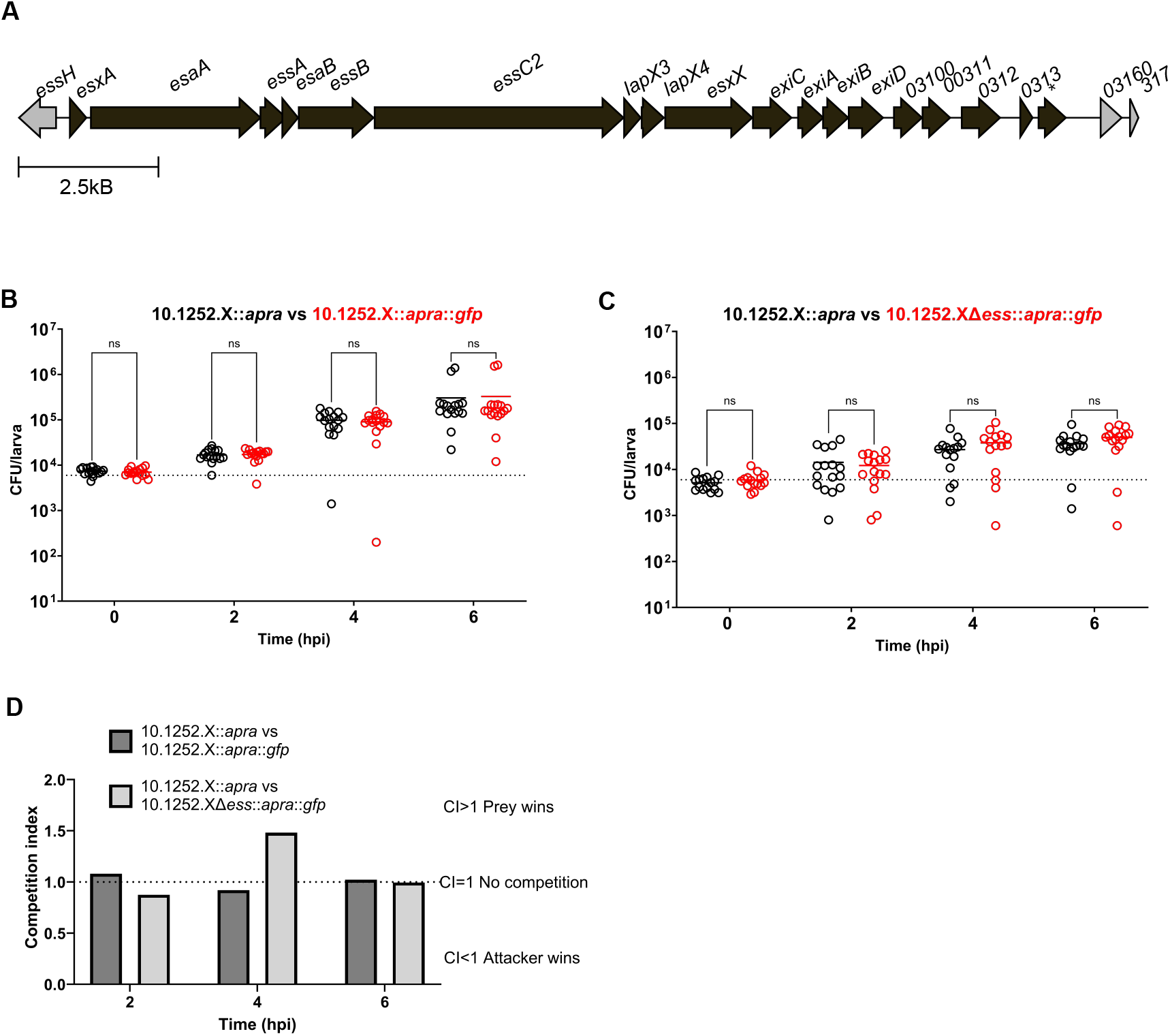
Assessing intrastrain competition in the zebrafish larval hindbrain. A. The T7SS/*ess* locus in 10.1252.X. Genes that are deleted in strain 10.1252.XΔ*ess* are shown in black. *indicates pseudogene. B and C. A 1:1 ratio of the indicated strains (each at 6000 CFU) was injected into the hindbrain ventricle of 3 dpf zebrafish larvae. B. 10.1252.X::*apra* (black) vs 10.1252.X::*apra*::*gfp* (red), n = 16 total larvae per timepoint, 2 biological replicates. C. 10.1252.X::*apra* (black) vs 10.1252.XΔessC::*apra*::*gfp* (red), n = 15-16 total larvae per timepoint, 2 biological replicates. At each timepoint larvae were sampled, homogenised and serial dilutions plated on TSA to determine CFU per larva. Circles represent individual larvae, horizontal bars represent the mean. Dotted lines indicate 6000 CFU (target dose). Multiple unpaired t-test with Welch correction. D. Competition indices at 2, 4 and 6 hpi.

## Discussion

The work described here adds to a growing body of evidence that the *S. aureus* T7SS is required for optimal virulence in animal models of infection, with a significant decrease in virulence associated with inactivation of the secretion system in multiple different strains and using different virulence models (e.g. (3, 4, 6, 8, 9)). However, it is not clear which secreted effectors, or combinations of effectors, are required for this process. Deletion of the gene encoding the nuclease toxin EsaD, or its chaperone, EsaE, from strain USA300 resulted in reduced IL-1β secretion in a murine bloodstream infection model, while deletion of *esaD* in strain Newman led to fewer abscesses and a reduced bacterial load in a similar model (26-28). By contrast, loss of EsaD in strain RN6390 did not significantly affect virulence in the zebrafish larval hindbrain model, despite the fact that EsaD was clearly active in this compartment because it was able to mediate killing of a competing *S. aureus* strain (10). Likewise, an *esxX* deletion mutant showed reduced abscess formation in murine skin and blood infection models, while we show no detectable loss of virulence in the zebrafish larval model for either *esxX* or *SAR0287* mutant strains (21). It should be noted, however, that using the same model, loss of another T7SS substrate, TspA, did show a decrease in zebrafish mortality, although this was non-significant (10).

Unlike EsaD and EsxX, which are only found in a subset of *S. aureus* strains, TspA is encoded by all *S. aureus* strains, and is therefore a prime candidate for a general T7SS effector that contributes to virulence. TslA is a second pan-*S. aureus* T7SS effector, which has phospholipase activity. While deletion of *tslA* did not result in a statistically significant difference in bacterial burden in a murine skin abscess model, the mutant strain did show a trend towards decreased virulence (9). It is possible that the reduced virulence seen across T7SS mutant strains results from the inability to secrete several effectors, each of which has a relatively small contribution to virulence.

The WXG100 protein and core secretion machinery component, EsxA, has been reported to modulate host cell apoptotic pathways in lung epithelial cells (7). As EsxA is no longer secreted when the secretion machinery is inactivated, the absence of extracellular EsxA could also partly account for the virulence defect of T7SS mutants. Alternatively, it is also possible that the decreased virulence may arise, at least in part, due to changes in the surface proteome of the bacteria either due to the loss of surface-localised T7SS substrates, or a change in physiology as a result of the secretion defect. Indeed, it was noted that inactivation of T7SS genes in RN6390 resulted in a transcriptional response that mimicked iron starvation, with changes in expression of several genes encoding cell envelope proteins (29, 30). It should also be considered that a combination of these factors may account for the changes in virulence, and that their contributions may also vary depending on the nature of the model and *S. aureus* strain used.

EsxX is toxic to *S. aureus* from both the cytoplasm and extracellularly, and *essC2* strains require two sets of immunity proteins that each provide compartment-specific protection (20). While this would suggest that the primary role of EsxX is bacterial antagonism, it should be noted that we were unable to demonstrate EsxX-dependent intrastrain competition *in vivo* in the zebrafish larval hindbrain. The conditions under which EsxX may deployed as an antibacterial toxin remain unclear.

## Conflicts of interest

The authors declare that there no conflicts of interest.

## Funding information

This study was supported by the Wellcome Trust through Investigator Awards 10183/Z/15/Z and 224151/Z/21/Z to TP. Work in the SM laboratory was supported by a Wellcome Trust Senior Research Fellowship (206444/Z/17/Z) and European Research Council Consolidator Grant (772853 - ENTRAPMENT).

## Author contributions

Conceptualization (TP, SM); Investigation (FU, GC, CM, AT, YY, MCG); Resources (GC, CM, JH); Visualisation (FA, FU); Writing – original draft (FA, TP); Supervision (TP and SM) and funding acquisition (TP, SM).

## Notes

### Competing Interest Statement

The authors have declared no competing interest.

