## Supplementary tables and figure for "The *Staphylococcus aureus* LXG-domain toxins EsxX and SAR0287 do not promote virulence in a zebrafish larval infection model"

**SUPPLEMENTARY INFORMATION**

**Table S1. Plasmids used in this study**

| Plasmid | Details | Reference/source |
|---|---|---|
| pBAD-18-Cm | Expression vector containing the inducible arabinose BAD promoter; CmlR | (1) |
| pBAD-SAR0287$_{LXG}$ | pBAD18-Cm encoding SAR_0287 N-terminal region (aa 1 - 314) | This work |
| pBAD-SAR0287$_{CT}$ | pBAD18-Cm encoding SAR_0287 C-terminal region (aa 315 – 556) | This work |
| pIMAY-apra-insertion | (*aac(3)-IVa*) under control of *rpsF* promoter, with flanking sequences from intergenic region downstream of *SAPIG0009* | This work |
| pIMAY | *E. coli/S. aureus* shuttle vector, temperature sensitive, CmlR | (2) |
| pIMAY-*esxX* | pIMAY carrying the flanking regions of *esxX* | This work |
| pIMAY-MRSAΔ0287 | pIMAY carrying the flanking regions of SAR0287 | This work |
| pIMAY-MRSA252Δ*ess* | pIMAY carrying the flanking regions of *essC3* | This work |
| pIMAY-GFP | pIMAY carrying the flanking regions between *SAPIG0102* and *SAPIG0103* to integrate *gfp* gene between them | This work |

**Table S2. Oligonucleotide primers used in this study**

| Primer | Sequence | construct |
|---|---|---|
| Bad_fwd | GGCATGCAAGCTTGGCTG | pBAD-SAR0287$_{LXG}$ |
| Bad_rev | TCTAGAGGATCCCCGGGTAC | pBAD-SAR0287$_{LXG}$ |
| SAR_0287 1-314_fwd | GTACCCGGGGATCCTCTAGAAGGAGGTTTCTAGTTATGGGGTACAAAGTTGATATG | pBAD-SAR0287$_{LXG}$ |
| SAR_0287 1-314_rev | AACAGCCAAGCTTGCATGCCTTATGCCATTTTTACTGCATTTTTTATTTTAATATTAC | pBAD-SAR0287$_{LXG}$ |
| CM18 | GCGCTCTAGACAGAGGAGGAGCCATGAGTGAATTTGCCCGTAATAATC | pBAD-SAR0287$_{CT}$ |
| CM19 | GCGCTCTAGACAGAGGAGGAGCCATGGGGTACAAAGTTGATATG | pBAD-SAR0287$_{CT}$ |
| pIMAY_fwd | TTGATATCGAATTCCTGCAG | pIMAY_ΔesxX |
| pIMAY_rev | ATCGATACCGTCGACCTC | pIMAY_ΔesxX |
| 305-up-500_fwd | TCGAGGTCGACGGTATCGATGATAAGGATGCTGATATAGC | pIMAY_ΔesxX |
| 305-up-500_rev | AAATCTCCTTTTTATACTCCTTTACTCTTTTATATTTATAATTG | pIMAY_ΔesxX |
| 305-down-500bp_fwd | GGGGTAATAAAAAGGAGATTTAAAATGAATAATACTAAG | pIMAY_ΔesxX |
| 305-down-500bp_rev | CTGCAGGAATTCGATATCAATAAACCAAAATGTGTTTTAGTTTTAC | pIMAY_ΔesxX |
| MGC394 | TATCGATAAGCTTGATATCGATTCATGGAATGCTTTAGAAG | pIMAY-apra-ins |

| | | |
|---|---|---|
| MGC401 | CTATAGGGCGAATTGGAGCTACCAAATGAAATACCAACAC | pIMAY-apra-ins |
| MGC311 | GCGCGAATTCATTCATGGAATGCTTTAG | pIMAY-apra-ins |
| MGC312 | GAAACTTTCCTCACTATTATACTTTT | pIMAY-apra-ins |
| MGC313 | TAATAGTGAGGAAAGTTTCAAATGAAT | pIMAY-apra-ins |
| MGC314 | GTATTGCACTTTATATTTGCACCTC | pIMAY-apra-ins |
| MGC315 | AAATATAAAGTGCAATACGAATGGCG | pIMAY-apra-ins |
| MGC316 | GTGCATTGGTCAGCCAATCGACTGGCG | pIMAY-apra-ins |
| MGC317 | ATTGGCTGACCAATGCACATAACAACA | pIMAY-apra-ins |
| MGC318 | GCGCGAGCTCACCAAATGAAATACCAAC | pIMAY-apra-ins |
| SAR287-A1 | CATGGAGCTCTCAATCAGCTCTATCTAATTATGAAAAC | pIMAY_ΔSAR0287 |
| SAR287-A2 | GGATAAGAAAGGGGCGTACCCCATAATAAATTTCCC | pIMAY_ΔSAR0287 |
| SAR287-B1 | ATGGGGTACGGATAAGAAAGGGGC | pIMAY_ΔSAR0287 |
| SAR287-B2 | GCGCGGTACCATATGATCTAACCAGCAATAAAT | pIMAY_ΔSAR0287 |
| SAR287-outfor | CCAAGCCTCTGTCAGCAAAG | pIMAY_ΔSAR0287 |
| SAR287-outrev | TTGGATATATATCTTTGTCCATG | pIMAY_ΔSAR0287 |
| SAR279-301-A | CATGGAATTCAATGTGCGTATACTGACCAC | pIMAY_ΔSAR0287 |
| SAR279-301-B | AGGTTTCTAGTTATGGCAATGAGCGACTTATCATAA | pIMAY_ΔSAR0287 |
| SAR279-301-C | AATATACGATGTTTATGATAAGTCGCTCATTGCCATAAC | pIMAY-ess-M252 |
| SAR279-301-D | CATGGAATTCCATAAAACGTTGTCTACTGG | pIMAY-ess-M252 |
| MGC402 | TATCGATAAGCTTGATATCGGACCTGACGTCGCTGCCG | pIMAY-GFP-ins |
| MGC403 | CAATCGCGATCCAAAAAGTCTTTAACACAAACAAAAAAGGAGGAAAC | pIMAY-GFP-ins |
| MGC404 | GACTTTTTGGATCGCGATTGCATGCCTG | pIMAY-GFP-ins |
| MGC405 | CTAGTTCATATATATCGCGAGCTGCATAAAAAAC | pIMAY-GFP-ins |
| MGC406 | TCGCGATATATATGAACTAGGGTGATTTAAAG | pIMAY-GFP-ins |
| MGC407 | TGGATCCCCCGGGCTGCAGGCAAAATTCGCATTTATAGCTAAAAATAATTTTG | pIMAY-GFP-ins |

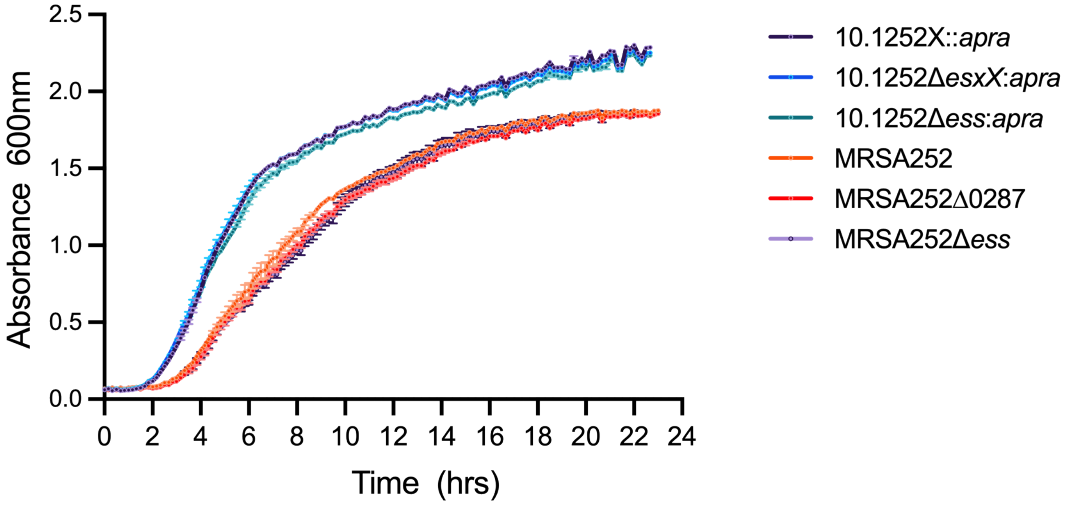


**Figure S1.** Growth curves for the indicated strains in TSB medium at 33°C.
